# Identifying Genes in Published Pathway Figure Images

**DOI:** 10.1101/379446

**Authors:** Anders Riutta, Kristina Hanspers, Alexander R. Pico

## Abstract

**BACKGROUND:** Pathway figures are commonly found in the biomedical literature providing intuitive models of complex processes in a visually concise format. The contents of a pathway figure often reflect the key findings and relevant context of an article. Unfortunately, the vast majority of pathway figures are drawn as one-off static images despite freely available pathway tools and resources, thus rendering their contents inaccessible to search, data mining and downstream analysis.

**APPROACH:** Leveraging advances in optical character recognition and domain expertise in pathway modeling, we devised an approach to identify genes in published pathway figures. The approach was optimized against a set of figure images obtained from PubMed Central and tested against a set of 400 curated pathways with known content from WikiPathways (F-measure 95.2%).

**RESULTS:** Applied to 3982 published pathway figures spanning a four year period, our approach identified 29,189 gene symbols representing 4159 unique gene identifiers. The gene content unlocked from just this small sample of published figures includes novel and diverse pathway associations unmatched by any pathway database. Our approach over doubled the number of genes associated with the articles containing these figures as compared to combined annotations available from PubMed and PubTator. Encouraged by these initial results, we plan to scale the approach to make the molecular contents of the continuing stream of published pathway figures more accessible.

## Introduction

Pathways are the grammar of biology. They describe the meaningful composition of molecules, interactions and reactions. In the biomedical literature, pathways are essential components often positioned as the first or last figure and used to convey the relevance and importance of a finding. We estimate that over 1000 pathway figures are published and indexed by PubMed Central (PMC) each month based on a sampling of PMC image search results. These figure images primarily consist of mechanistic pathway diagrams drawn using general-purpose illustration software, but also include a minority of interaction network representations generated using graph modeling tools. Only a tiny fraction are properly annotated models sourced from a public database. Thus, the vast majority of these figures are drawn as one-off creations, disconnected from visualization and annotation standards, and then exported as static image files, rendering their contents invisible to search, data mining and bioinformatic analysis. Important information summarizing the context and novel models in biomedical research in the form of pathway figures is practically inaccessible to further discovery, reproducibility and meta analysis efforts.

The general problem of extracting information from the biomedical literature is an established and active area of research involving the processing of abstracts, full texts, captions and images (1–4). These methods have made their way into popular annotation services, such as PubTator, which mines the abstracts of all PubMed articles for genes, diseases, species, chemicals and mutations, and also makes these results available in a searchable database (www.ncbi.nlm.nih.gov/CBBresearch/Lu/Demo/PubTator) (5). Figures, however, are more challenging than text documents. The majority of approaches have targeted domain-specific figures (3), such as bar charts (6), axis diagrams (7), phenotype images (8–10) and gel electrophoresis images (11); while others have targeted heterogeneous figures, such as Yale Image Finder (12) and FigureSearch (13–15). Of the approaches that handle pathway figures, a few were available as public resources for a time but are now obsolete, including BibText (16), Figurome (17) and BiologicalNetworks (18). There is no resource that provides updated access to the gene content locked away in the stream of pathway figure images being published.

Approaching this problem from the perspective of pathway database maintainers, we are particularly focused on the large and growing gap between the knowledge expressed in pathway figures and the information captured in annotated pathway databases. We started the WikiPathways project (www.wikipathways.org) a decade ago in an attempt to address this gap, relying on communities of researchers to model the full diversity of pathways in coordination with discovery and publication (19). While this approach has been wildly successful in contrast to traditional, internal-only, canonical pathway databases, the combined efforts of all pathway databases (including WikiPathways) fail to capture newly published information at a rate or scale even close to real time. The long-term goal of the work presented here is thus to enrich pathway databases with content that has eluded prior efforts, initially at the resolution of basic gene sets to enable search, associations and enrichment analyses. The identities and relative positions of genes in pathway figures can inform modeling efforts to capture interactions and other higher resolution content. The genes identified in pathway figures can also feed into gene-article association databases (e.g., PubTator), general gene set enrichment resources, and article lookup services. We are dedicated to making our code, methods and results freely and openly available per FAIR principles (20).

The aim of this study is to assess the gene content in pathway figures, relying on a publicly available source of published figures, gene nomenclature, optical character recognition (OCR) technology and heuristics derived from our experience as pathway biology researchers and biocurators (Figure 1). We describe the technical development and performance of our Pathway Figure OCR (PFOCR) method and apply it to a sample of the biomedical literature. We find a veritable cornucopia of genes collected in nuanced pathway representations far exceeding current pathway databases and gene-article association resources. These results motivate continued development and broader application of the method toward the goal of making pathway information more accessible and useful to researchers.

**Fig. 1.**
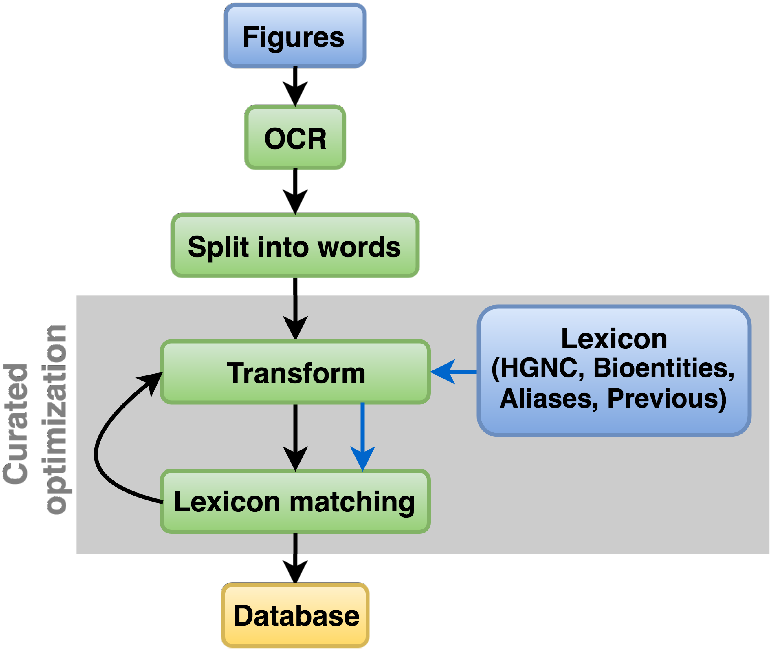
Pathway Figure OCR (PFOCR) Method Overview. The inputs (blue) include pathway figure images downloaded from PMC and lexicon sources. Processing steps are represented in green. Results at multiple stages were stored in a PostgreSQL database (yellow). The Curated Optimization phase of the method development involved iterations within the shaded region.

## Methods

### Pathway figure collection

A PubMed Central image search using the query “signaling pathway” generated 37,032 results (April 2015). Displaying 100 results per page, the HTML content for the first 5000 results were downloaded and then parsed by a PHP script to collect JPG figure image files and metadata on figure number, title, caption and URL. Visual inspection of these 5000 image files from publications spanning 2011–2015, revealed that 3982 (79.6%) actually contained a pathway diagram or network; the remainder consisted of various tables and gel electrophoresis images that happened to contain the words “signaling” and “pathway” in their captions.

### Optical character recognition

The 3982 signaling pathway figure image files were submitted to Google Cloud Vision (GCV) API for “text detection” by OCR (cloud.google.com/vision/docs/detecting-text). An input JSON payload was prepared including the image as a base-64 text string and a specification for “text detection”. The service returned a JSON response per image containing a list of recognized words split by newline and space. A Python script managed the API submission and results handling, as well as the postprocessing and lexicon matching. A PostgreSQL database stored the OCR results, along with the lexicon, intermediate post-processing results, matches and summaries for each run of the pathway figure OCR system. The scripts and table schemas can be found in the project code repository (see Data Availability).

### Lexicon construction

Our lexicon combined four types of human gene symbols from two sources mapped to NCBI Gene identifiers: HGNC (symbol, previous and alias) (www.genenames.org), and Bioentities (relations) (github.com/wikipathways/bioentities forked from github.com/johnbachman/famplex). Symbols and corresponding identifiers were downloaded in tabular formats from these sources and processed in Excel following a protocol specified in the lexicon directory in the project code repository, resulting in a set of 4 CSV files (see Data Availability). Given the overlap between these sources, they were loaded into a combined database table in preferential order based on our experience with symbol usage by pathway figure authors and biased against false positive hits: HGNC symbol > Bioentities > HGNC alias > HGNC previous. Thus, if the same symbol from HGNC symbol and HGNC alias maps to different NCBI Gene identifiers, then we only included the HGNC symbol mapping in our lexicon and discarded the conflicting entry from the lesser source. The final lexicon used in this study, following Curated Optimization, included 58,242 unique symbols mapping to 19,176 unique identifiers.

### Post-processing

The words isolated from the OCR results included gene symbols, aliases and common names, in addition to references to sets of genes (e.g., “IRF1-3”) and other labels and text embedded in the image. A series of transformations were applied to these words in order to improve matching with our lexicon. Transformations applied to both the OCR-extracted words and the lexicon entries are called *normalizations*, while the transformations applied only to the OCR-extracted words are called *mutations*. A total of eight transformations were developed and applied in this study (Table 1). The order and combination of transformations had a significant effect on the results. The order in the table essentially matches the algorithm settled upon during the Curated Optimization. The precise algorithm is available in `match.py` and the `transforms` directory in the code repository (see Data Availability).

**Table 1.** Transformations used for matching OCR-extracted words to lexicon entries. The set of eight transformations including “mutations” applied only to words and “normalizations” applied to both words and lexicon. Each transformation has a general description and one or more specific examples listed.

| Name | Description | Type | Examples |
| --- | --- | --- | --- |
| **upper** | Force upper case | normalization | Akt1 => AKT1 |
| **stop** | Exclude due to high false positive occurrence, ambiguity or lack of specificity | normalization | NOT, FOR, TYPE<br>S21, CYCLIN |
| **nfkc** | Convert characters to compatible forms | normalization | TP<sup>53</sup> => TP53 |
| **deburr** | Remove combining characters | normalization | ĆĎÇ2 => CDC2 |
| **expand** | Extract gene names from conjunctions, enumerations and ranges | mutation | IRF1,3-4 => IRF1, IRF3, IRF4<br>YAP/TAZ => YAP, TAZ |
| **root** | Remove prefixes and suffixes | mutation | AKT1-p, GST-AKT1, WNTs |
| **swaps** | Substitute Greek symbols, non-standard names and commonly mistaken characters in specific contexts | normalization | α, alpha => A<br>III => 3<br>P13K => PI3K |
| **alphanumeric** | Remove non-number/letter characters | normalization | MMP-9 => MMP9 |

### Curated optimization

In order to address false positive and negative matches in an intelligent way, an iterative process of curated optimization was developed and performed (Figure 1). After running our PFOCR system on the sample set of 3982 signaling pathway figures, 30 figures and their transformed lexicon-matching results were randomly selected and reviewed by experienced pathway biocurators, confirming true positives and characterizing false positives and false negatives. These observations were used in the design and refinement of our lexicon and transformations, prioritizing the most common and tractable errors. The same 30 pathway figures were assessed in a run following the updates to verify intended fixes. Then another set of 30 random pathway figures and results were selected to initiate another round of curated optimization. This process was repeated for five consecutive rounds, resulting in a dramatic drop in the occurrence of tractable improvements (see Supplemental Figure S1).

### Performance measures

In order to assess the performance of our optimized system, it was applied to a distinct set of 400 PNG images corresponding to human pathway models from WikiPathways (downloaded on 04/17/2018). Since the WikiPathways models are annotated with gene identifiers, performance could be precisely quantified based on true positive, false positive and false negative counts (Table 2 and Figure 2). JSON payloads containing a pointer to the image and associated metadata were prepared as inputs for OCR by GCV and the results were processed and represented in the same manner as those from published pathway figure images. A set of R scripts were developed to compare the identified gene sets against the known gene contents per pathway and to calculate standard performance measures (see Data Availability for `performance` directory in code repository). The lexicon was used to restrict the space of possible matches. The automatic expansion of hits provided by the inclusion of Bioentities in the lexicon were manually confirmed and counted as true positives rather than false positives since they were adding legitimate content consistent with the biology represented in pathways.

**Table 2.** Performance on Pathways with Known Gene Content. Standard performance measures were calculated based on a test set of 420 annotated human pathway models from WikiPathways with known true positive, false positive and false negative counts per pathway. F-measure is the harmonic average of precision and recall. The medians and standard deviations are presented in the table.

| Precision | Recall | F-measure |
| --- | --- | --- |
| 100% (1.9%) | 90.9% (11.4%) | 95.2% (13.4%) |

**Fig. 2.**
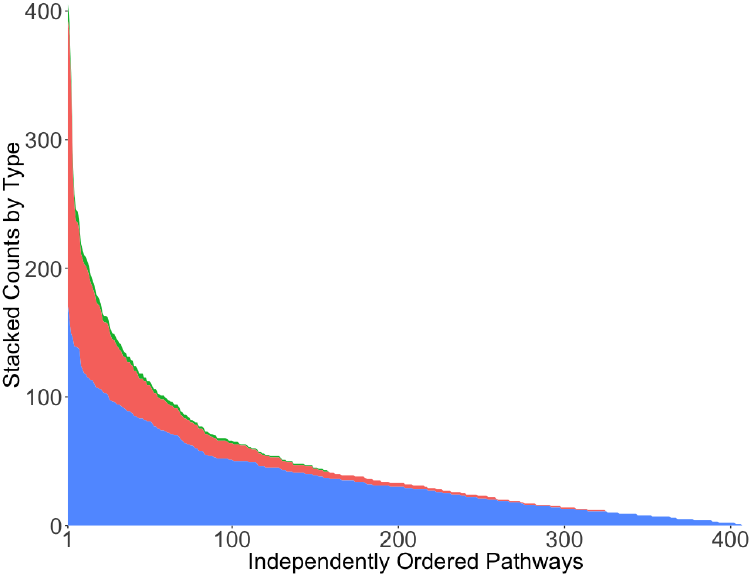
PFOCR performance on WikiPathways test set of 420 pathways for Homo sapiens. Provided the known gene content in these previously curated pathway models, we calculated true positive (blue), false negative(red) and false positive (green) counts. Independently ordered by each count, the plot of stacked areas shows very low false positive counts and a favorable balance of true positive vs false negative with a few exceptions (5%) where more genes were missed than detected. For an alternative ordering by pathway, see Supplemental Figure S2.

## Results

### Assessing the PFOCR Method

We have developed a methodology to proficiently identify gene symbols extracted from a diverse set of published pathway figure images. A sample set of 3982 figures was collected from a PubMed Central image search result and pruned by visual inspection. Words were extracted from the images by OCR using Google Cloud Vision and matched to a lexicon of human gene symbols and corresponding NCBI Gene identifiers through a series of transformations. We optimized the method by manually inspecting randomly selected figures over multiple rounds, incrementally curating the lexicon and set of transformations. This Curated Optimization was conducted with an explicit bias against false positive results to avoid associating irrelevant and inaccurate genes with figures and papers. Using WikiPathways (21) as a source of pathway figures with independently preidentified content, we were able to calculate the number of true positive, false positive and false negative gene identifications over a larger set of figures with greater accuracy and precision than by manual inspection. The median true positive count in the WikiPathways set is 28 genes per pathway. The median false negative count is three genes and median false positive count is zero (Figure 2). The median F-measure across these curated pathway figures is 95.2% (Table 2). These performance measures reflect our bias against making false gene identifications at the cost of non-negligible numbers of missed gene identifications. On a per pathway basis, we observed that a fourth of the false negative cases were due to poor color choices (i.e., text/background contrast) and another tenth were due to using non-standard gene names.

Overall, given the simplicity of the approach using publicly available resources, the performance of the PFOCR method is striking. Note that the method was not optimized on the WikiPathways content, but rather on a wildly diverse set of signaling pathway images. Granted, with a few exceptions, the WikiPathways content tends to be clearer than the typical pathway figure due to the guidance, standards and community curation provided. Thus, we do not claim, nor expect, the same performance measures to extend to the full diversity of pathway figures found in the literature. Furthermore, the method will require further rounds of optimization when applied to other species and to processes other than signaling, such as metabolic and regulatory pathways; aspects of the transformations (e.g., swaps) and lexicon (e.g., Bioentities) are biased for dealing with signaling genes and human gene nomenclature. Nevertheless, based on the performance observed during Curated Optimization, the potential for further optimization and the calculated recall and precision under above-average conditions, it is clear that the technology and methods presented here are adequate for interrogating published pathway figures for relevant gene content.

### Applying the PFOCR Method

When we approached this problem in early 2015, the public beta release of GCV was still eleven months away and Google’s Tesseract and Adobe’s Acrobat Text Recognition were two popular, affordable options. Table 3 presents the results from the 2015 run of Acrobat and Tesseract in comparison with GCV results from 2018, using either the 2015 lexicon and post-processing (second row; see asterisk) or our latest methods presented here (top row). The same set of 3982 signaling pathway figure images were used in all cases. Our optimized method together with GCV identified a total of 29,189 gene symbols, a median of 15 genes per pathway figure, which map to 4159 unique gene identifiers. Intriguingly, 1461 of these unique genes are not present on any pathway in the current WikiPathways database, thus representing novel pathway information. Advances in OCR technology and in processing each improved the detection of novel pathway information by 90% relative to the current collection of WikiPathways models. In terms of coverage, we identified 66.2% of the lexicon-bounded gene content present in the subset of 120 curated human signaling pathways from the WikiPathways database. Recall, this is from a relatively small sample of signaling pathway figures published over just a 4-year period. In the next section, we attempt to extrapolate these findings to determine how many pathway figures would need to be processed before redundancy outweighs novelty in terms of gene content.

**Table 3.** Improvements in OCR and Post-processing Results. “Hits” refer to words found in the OCR results that match an entry in our lexicon through transformation. “IDs” refer to NCBI Gene IDs. Because we included HGNC aliases and previous symbols in our lexicon, some IDs correspond to multiple hits, and because we included Bioentities in our lexicon, some hits correspond to multiple IDs.

| Tool | Figures | Total Hits | Unique Hits | Unique Gene IDs | Novel Gene IDs |
| --- | --- | --- | --- | --- | --- |
| **GCV(2018)** | **3982** | **29,189**<br>(+46%) | **4865**<br>(+23%) | **4159**<br>(+57%) | **1461**<br>(+90%) |
| GCV* (2018) | 3982 | 19,933<br>(+53%) | 3867<br>(+69%) | 2657<br>(+89%) | 767<br>(+89%) |
| Acrobat (2015) | 3982 | 12,645 | 2366 | 1471 | 404 |
| Tesseract (2015) | 3982 | 13,490 | 2199 | 1337 | 409 |

### Assessing the Gene Content in Pathway Figures

A plot of accumulated unique genes as a function of processed pathway figures shows a clear power relationship described nicely by a square root function (Figure 3). Given the restricted search space (i.e., published signaling pathway figures), we would expect a plateau as the total number of genes in such figures is approached. We have not collected enough data yet to estimate this number, but the beginning of a plateau may be evident toward the end of the plot. Just prior to the plateau, 4000 unique genes were collected after processing 2925 images. By contrast, 269 randomly selected human pathway models from WikiPathways were sufficient to collect 4000 unique genes using the same OCR processing methods. The estimated 10-fold decrease in collection efficiency from figure images is due to a combination of there being fewer detected genes per image (median of 15 genes per figure image versus 31 genes per WikiPathways image) and greater redundancy across figure images versus a curated database of pathway models (see Figure 4). In an unrestrained search space, the model predicts that we would need to process a total of 12,000 pathway figure images to double our current collection to total 8000 unique human genes. By monitoring such a plot one can readily evaluate diminishing returns in any future image collection and processing effort. Expanding the search space (e.g., to metabolic pathways, etc) would shift the total unique gene count limit and allow more efficient collection until, of course, the absolute number of unique genes in published figures is approached. A continued stream of new discovery, including new roles and contexts for previously captured genes will motivate continued figure processing even after every gene is captured at least once.

**Fig. 3.**
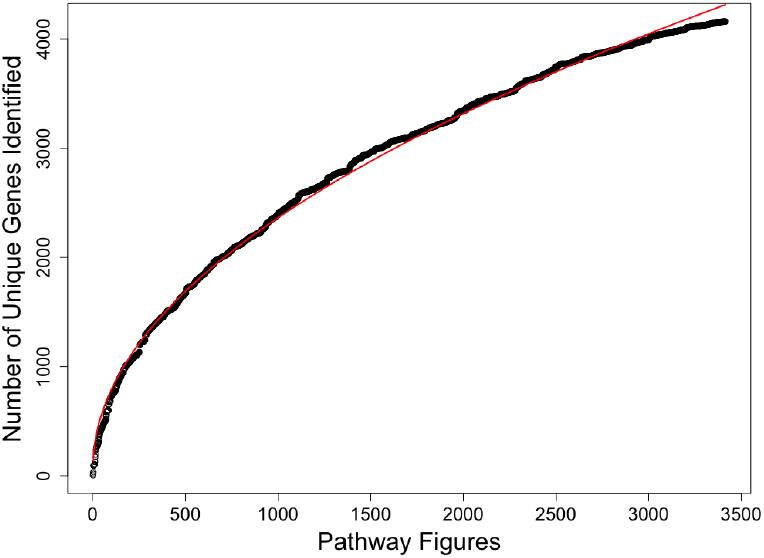
Number of accumulating unique genes detected as a function of OCR-processed pathway figure images. The unique gene content across the set of 3982 signaling pathway figures is plotted as an accumulating count (black circles). A square root function (red line) describes the data(R2=0.9971). Deviation from the line is observed toward the end of the plot, perhaps indicating diminishing returns as the total number of unique genes in the search space is approached.

From another perspective, the annotated clusters in Figure 4 represent the diversity and coherence observed in complete results (Supplemental Figure S3). We find easily-identified canonical pathways with only minor variation (e.g., mTOR signaling cluster), as well as highly variant sets of pathways that defy traditional classification (e.g., Misc. signaling cluster). These results illustrate important characteristics of the pathway content represented in these 3982 images. First, while there are a few common pathways present at high redundancy (e.g., TLR signaling), the content overall is quite diverse. No two pathway images in this collection contain the exact same set of genes. This is relevant to their potential use as a source of gene sets for enrichment analysis, where uniqueness is required to allow ranking. Second, the identification of clusters within this diversity suggests the potential for annotating publications with pathway names and links to other publications or pathway resources based on cluster assignment, i.e., distance to pre-annotated clusters.

**Fig. 4.**
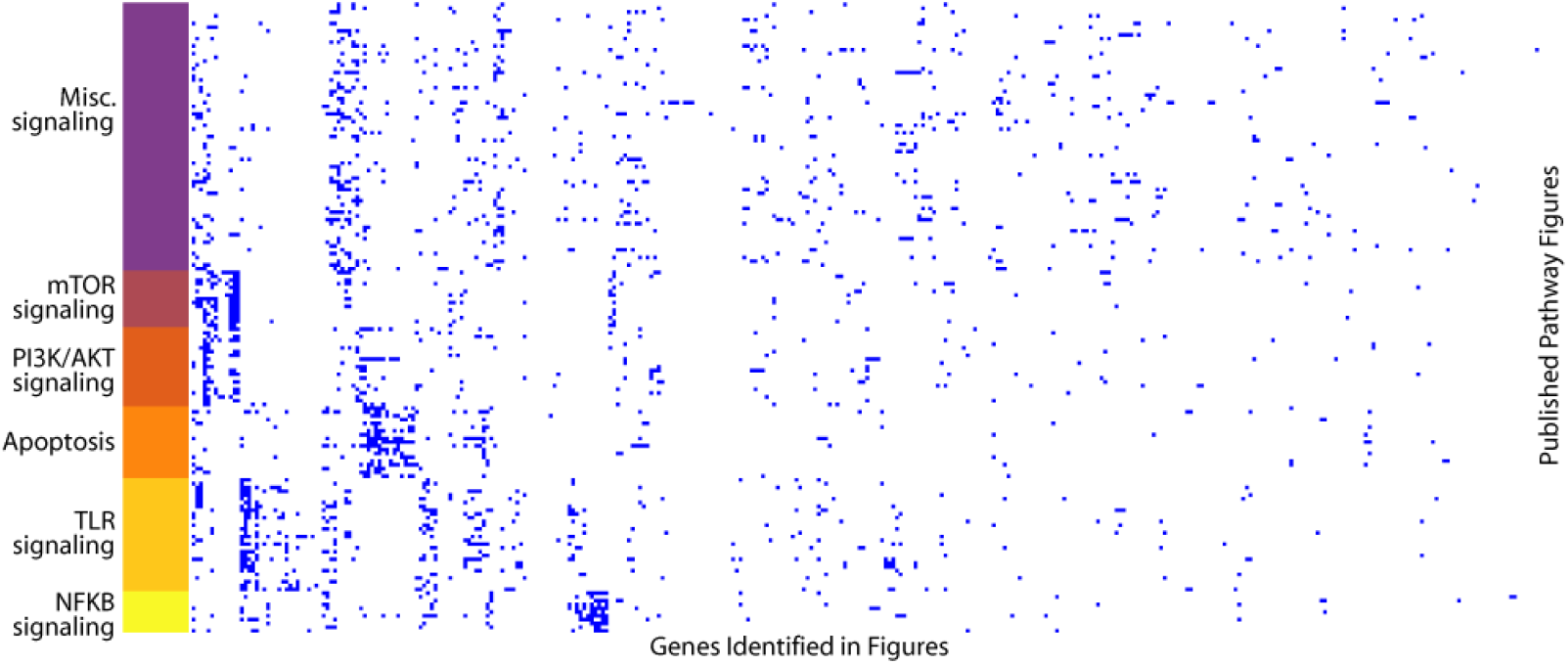
Six representative clusters of published pathway figures based on identified genes. Rows are PMCIDs and columns are NCBI Gene IDs. The complete clustering results are shown in Supplemental Figure S3. Clusters 8-13 are shown here (same color coding) and annotated based on manual inspection. The annotation roughly corresponds to well-known pathway categories with the exception of the miscellaneous cluster.

From yet another perspective, the PFOCR system can be used to annotate articles with gene identifiers. Existing annotations available from PubMed and PubTator provide genes identified from from abstracts and full text. For the subset of articles with pathway figures, however, we can significantly augment these gene annotations (Figure 5). Limited to the same lexicon and set of articles, we found that the PFOCR method identified 75% of the genes previously identified from the text by PubMed and PubTator, suggesting that the pathway figures themselves capture a significant proportion of the paper’s findings in terms of gene content. An additional 2603 genes were identified by PFOCR that were not found by these other text-based approaches, representing a 128% increase in gene annotations for this sample of articles.

**Fig. 5.**
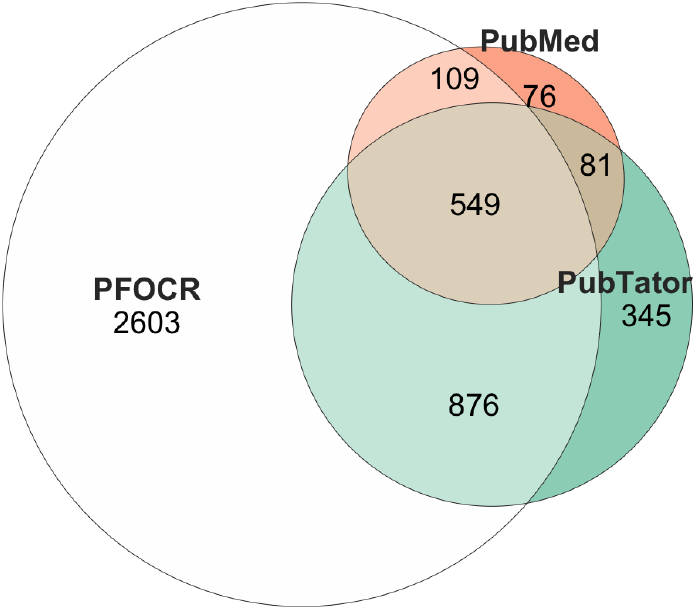
Unique genes with counts as identified by PFOCR, PubMed and PubTator. The numbers of unique genes identified by our PFOCR method, PubMed and PubTator over the same lexicon and set of articles (excluding 58 articles lacking PMIDs). The PFOCR method identified 75% of the genes extracted from the text by PubMed and PubTator, and an additional 2603 genes not found in either of these text-based approaches.

### Limitations and Future Work

The most apparent limitation of this study is the focus on a relatively small sample of 3982 signaling pathway figures published over a 4 year period. Based on PMC queries for figures titled or captioned with “signaling pathways”, we observe 82,179 image results spanning January 2010 to January 2018. Extrapolating the same proportion of *actual* pathway figures we observed in our sample of 5000 images (79.6%), we estimate there are 65,000 figure images of signaling pathways. The rate has increased year-to-year over this time period, reaching just over 1000 signaling pathway figures per month in recent years, based on our estimated proportion. PMC image queries for “pathways” returns approximate 4 times as many results, though we suspect the proportion of actual pathway figures in these results to be much lower. We intend to apply the PFOCR method to an ongoing collection of newly published pathway figures, backfilling to some date to be specified based on redundancy and utility of the extracted gene content. This system will involve replacing our manual inspection of images with automated strategies that can either positively identify actual pathway figures (e.g., by learning features of captions) or filter out identifiable figures of other types, e.g., gel electrophoresis (11).

In addition to extending the scale of processed images over time and search space, we also intend to expand the scope of organisms and molecular types (e.g., miRNA, drugs and metabolites). This will involve expanding and diversifying the lexicon and transformations. The Curated Optimization approach worked well at the scale of this study, but may become too time intensive and inefficient for a broader scoped endeavor. We are pursuing collaborations in the NLP, named entity recognition and image processing fields already specializing in biomolecular content and research literature.

Decades of prior experience annotating, curating and using biological pathways were applied to the rule-based design of our lexicon and transformations. Our experience was a key factor in our decision *not* to use machine learning, relying instead on the pattern matching and intuition we have organically acquired. This type of learning is never complete. We already have a backlog of improvements to apply to future work, including refining word splitting and handling homoglyphs. One of the updates will be to include *weighted* and *fuzzy* lexicon matches. As we build a corpus of gene sets per figure, we could use prior observations to help weight two or more possible matches using, for example, a guilt-by-association score. Weights could also be informed by whether the match is also mentioned in the caption or text of the article, which lexicon source it is matching, how many transformations were applied, how many confusable characters the match contains (unicode.org/cldr/utility/confusables.jsp), etc.

In terms of the OCR component, we anticipate continued improvements to services that are already impressive and affordable. Alternatives to GCV are now available from Amazon (aws.amazon.com/rekognition), Microsoft (www.microsoft.com/en-us/research/research-area/computer-vision) and others. Regardless of provider, there are optimizations to the images that can be made prior to feeding it in, such as greyscale, contrast enhancement, segmentation and rotating for mixed text orientations.

The greatest limitation by far, however, is the low standard of pathway figure preparation and review in the published literature. Not only is there a lack of standards in place for pathway figures, but basic data visualization practices and even legibility are also often neglected. Numerous pathway drawing tools and resources are available to researchers, for free, including WikiPathways (21), Reactome (22), SBGN (23) and SBML editor (24). Any of these would provide significant improvements to multiple aspects of current pathway figures. The pathways from such tools and databases conform to standards relating to gene identification, labeling and depiction that will improve OCR performance (see benchmarking against WikiPathways). Furthermore, if a proper pathway modeling tool is used, then the need to extract genes post-publication is resolved upfront. Databases like Reactome and WikiPathways can accommodate pathway models needed for any publication or presentation purpose; they simultaneously maintain a richly annotated model to support search, bioinformatic analysis and data visualization.

## Conclusions

A wealth of pathway information published only as figures is lost to further investigation by programmatic and sometimes even manual means. By applying off-the-shelf OCR, a simple lexicon and optimized set of heuristic transformations, we have assessed the quantity, quality and diversity of genes identified in pathway figures. From a sample set of 3982 signaling pathway figures published over a four year period, we identified a total of 29,189 gene symbols, a median of 15 genes per figure, which map to 4159 unique gene identifiers. Over a third of these genes were not previously associated with any pathway in the WikiPathways database. Clustering the genes identified in figure images revealed a diverse mix of both canonical and nuanced pathways, with no two pathways being identical. This poses a meaningful challenge to the dogmatic practice of current pathway databases to curate one “true” version of a process, such as apoptosis, that in actual usage has quite variable representations.

This study was limited to a relatively small sample of signaling pathway figures and to the identification of only genes, ignoring for now miRNA, drugs, metabolites and process terms. We intend to continue the development and broaden the application of our method in future work. Nonetheless, the gene content alone has great practical value for indexing articles, making gene associations and supporting gene set enrichment. Providing these data in public search tools and resources will greatly enhance the scope and scale of pathway analysis.

Pathway data, and more generically interaction data, are among the many data types effectively neglected in our current publishing and knowledge accumulation practices. Eventually the current trend toward proper data modeling and deposition in scientific publishing will be standardized, e.g., as a requirement for funding. This change will make the *post hoc* extraction of biological information from future text, PDFs and images less urgent. In the meantime, knowledge extraction efforts are critical to keeping up-to-date with the publication deluge, accessing older publications and making the contents of articles accessible to search, data mining and bioinformatic analysis. Furthermore, for people with visual impairments, published figures in general are often not accessible (25). The National Library of Medicine even admits, “Some parts of the PubMed Central archive may not be fully accessible to persons with disabilities, even with assistive technology.” (www.ncbi.nlm.nih.gov/pmc/about/accessibility). By transforming images into annotated resources this work opens up possibilities for making pathway figures accessible to all.

## Data Availability

Code Repository: github.com/wikipathways/pathway-figure-ocr

## Supplemental Figures

**Fig. S1.**
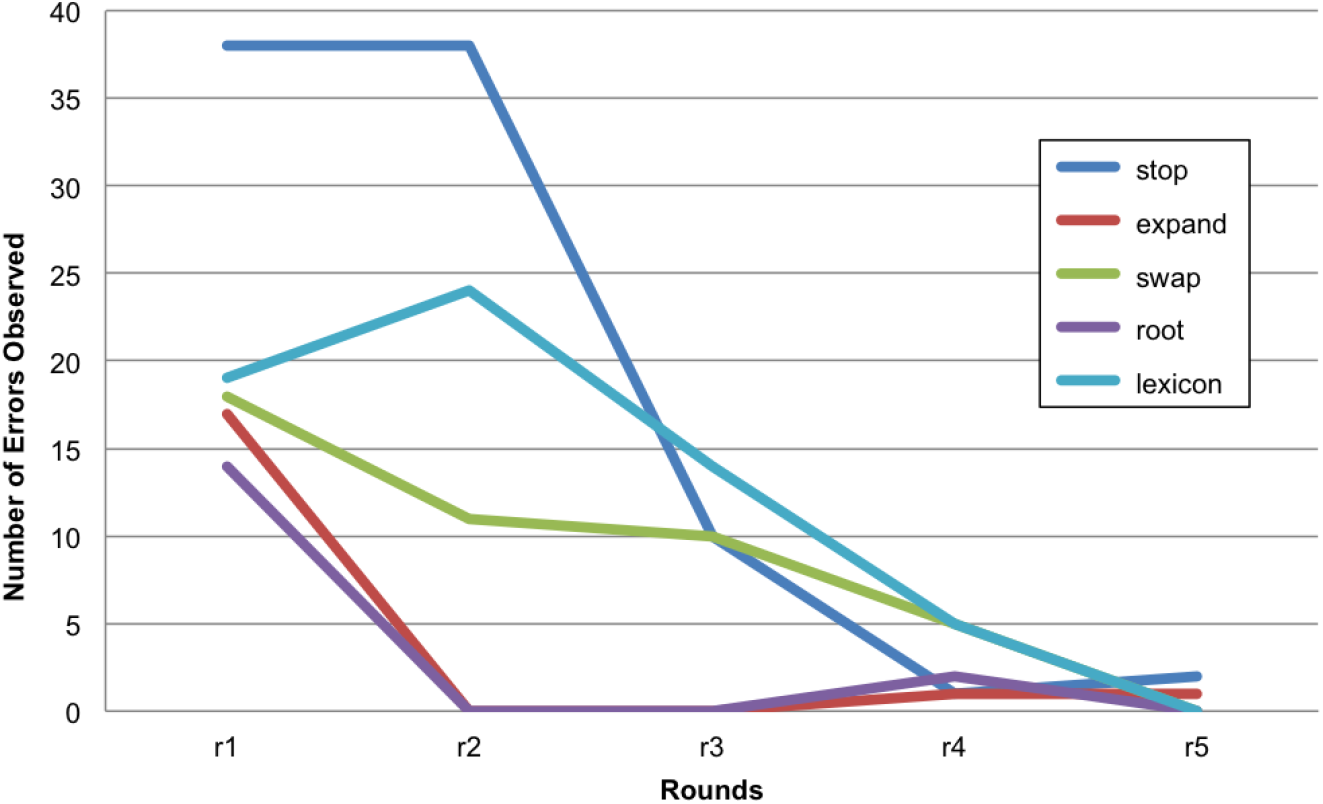
Number of Observed Errors by Optimization Type. Errors addressable either by updating transformations or the lexicon were counted over five successive rounds of OCR and processing (r1-r5) in order to direct the curated optimization of the PFOCR method is a way that was directed, efficient and biased against false positives. The error types are labeled by the transformations that were being refined to address them (e.g., stop, expand, swap, root; see Table 1) or labeled as lexicon in cases where the lexicon needed refinement.

**Fig. S2.**
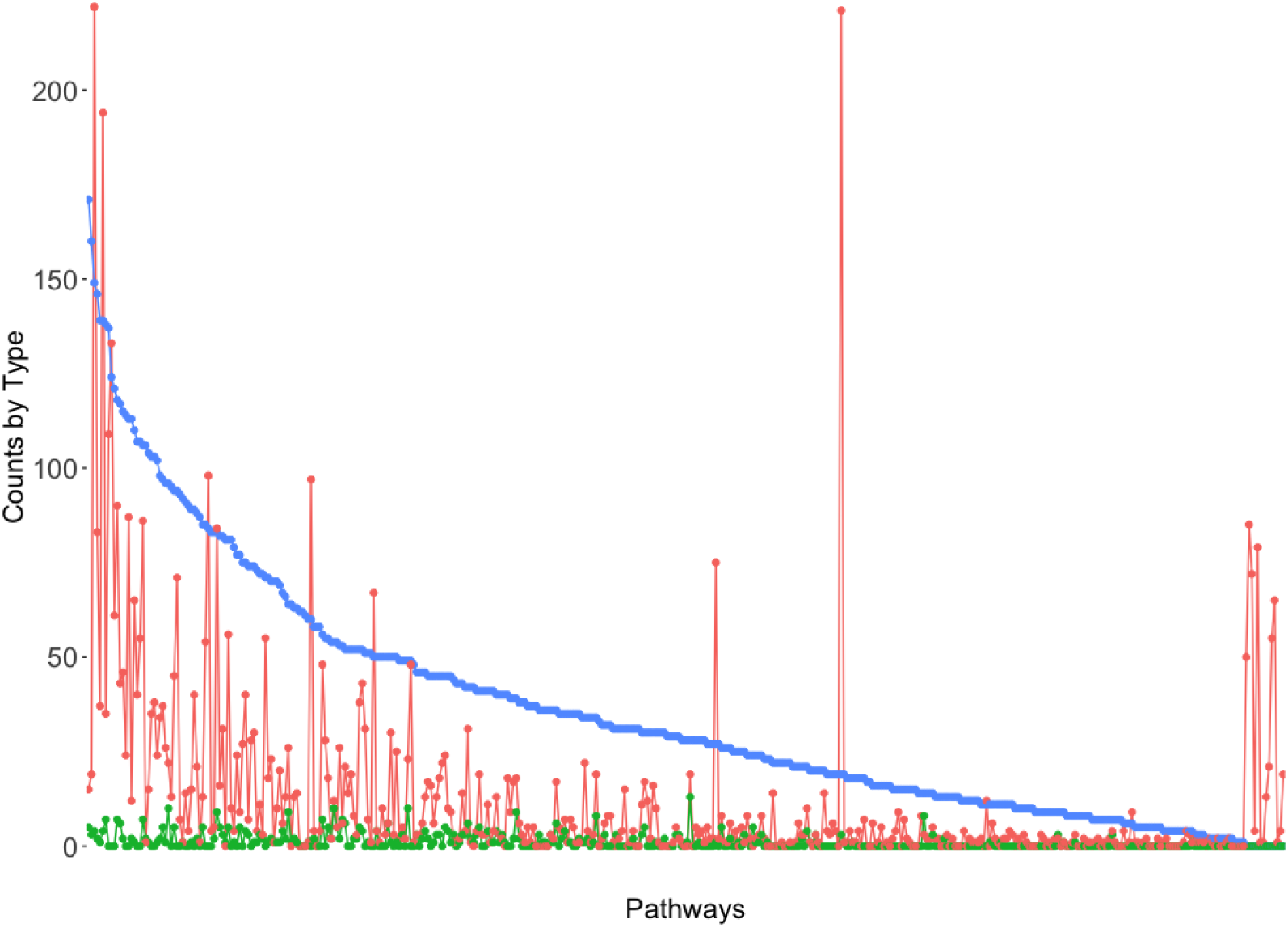
PFOCR performance on WikiPathways sample set of 420 pathways for *Homo sapiens*. Provided the known gene content in these previously curated pathway models, we calculated true positive (blue), false positive (green) and false negative (red) counts. Ordered by true positives per pathway, the plot shows very low false positive counts and a favorable balance of true positives vs false negatives with a few exceptions (red spikes, 5%) where more genes were missed than detected. Automatic expansion of Bioentities hits were manually confirmed and classified as true positives rather than false positives since they were adding legitimate content.

**Fig. S3.**
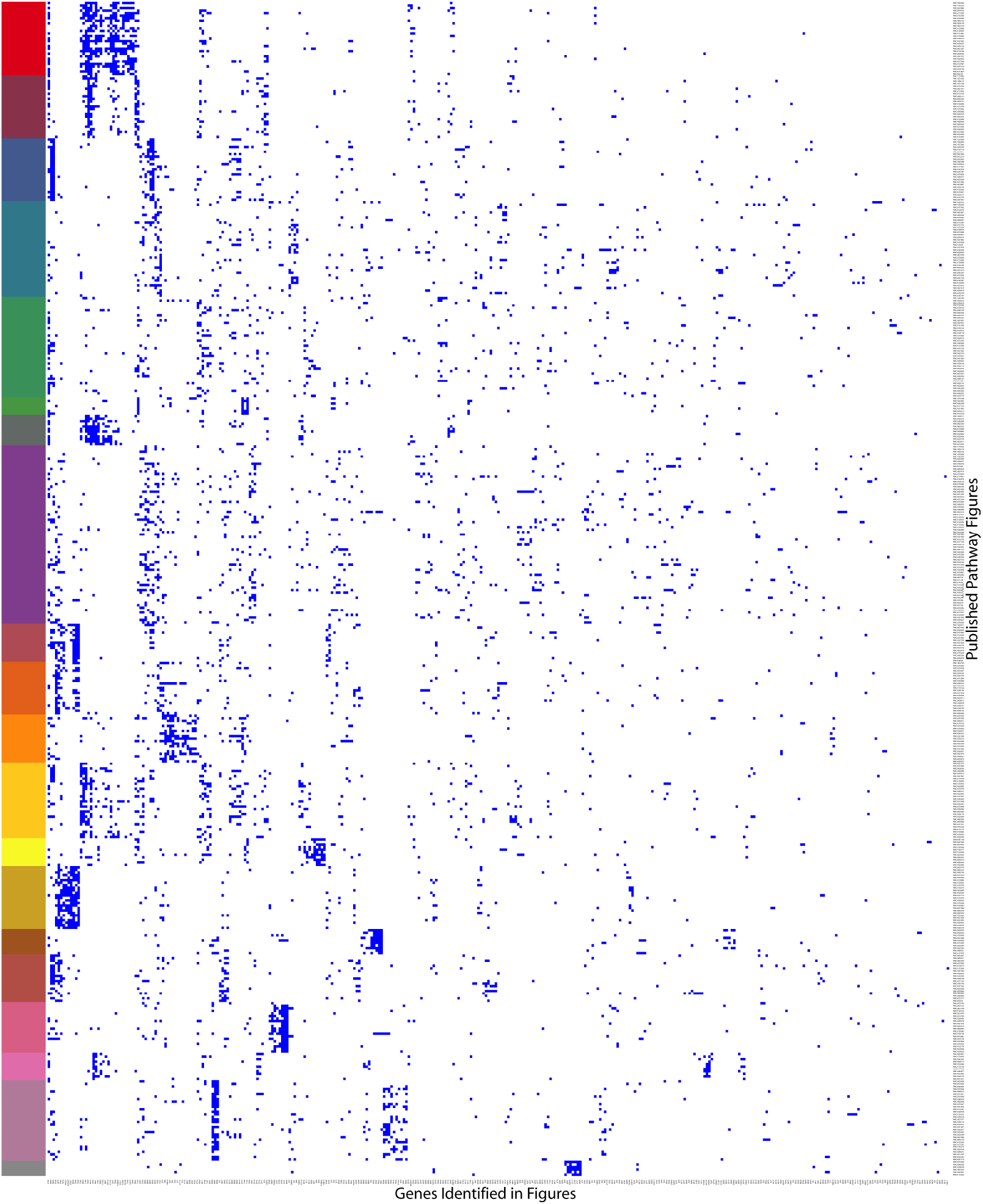
Binary heatmap of hierarchical clustering of genes identified in published pathway figures, excluding those inferred by Bioentities. Rows are PMCIDs and columns are NCBI Gene IDs. Clustering was performed on a subset of data, first excluding all genes matching and inferred by Bioentities and then only the most frequent genes occurring in at least 11 images (363 NCBI Gene IDs) and figure images containing at least 8 of these genes (466 PMCIDs). The detection of a gene on a given pathway is indicated in blue. An arbitrary cutoff of 20 clusters produced the row color strip and was used to annotate by pathway names where possible.

